# Quantitative proteomics reveals tissue-specific, infection-induced and species-specific neutrophil protein signatures

**DOI:** 10.1101/2023.09.05.556395

**Authors:** Gabriel Sollberger, Alejandro J. Brenes, Jordan Warner, J. Simon C. Arthur, Andrew J.M. Howden

**Author notes:** Corresponding authors: Gabriel Sollberger and Andrew Howden.

## Abstract

Neutrophils are one of the first responders to infection and are a key component of the innate immune system through their ability to phagocytose and kill invading pathogens, secrete antimicrobial molecules and produce extracellular traps. Neutrophils are produced in the bone marrow, circulate within the blood and upon immune challenge migrate to the site of infection. We wanted to understand whether this transition shapes the mouse neutrophil protein landscape, how the mouse neutrophil proteome is impacted by systemic infection and perform a comparative analysis of human and mouse neutrophils. Using quantitative mass spectrometry we reveal tissue-specific, infection-induced and species-specific neutrophil protein signatures. We show a high degree of proteomic conservation between mouse bone marrow, blood and peritoneal neutrophils, but also identify key differences in the molecules that these cells express for sensing and responding to their environment. Systemic infection triggers a change in the bone marrow neutrophil population with considerable impact on the core machinery for protein synthesis and DNA replication along with environmental sensors. We also reveal profound differences in mouse and human blood neutrophils, particularly their granule contents. Our proteomics data provides a valuable resource for understanding neutrophil function and phenotypes across species and model systems.

## Introduction

Neutrophils are one of the most abundant cell types in the immune system. Together with eosinophils and basophils, they belong to the family of granulocytes, which develop in the bone marrow and circulate in the bloodstream to patrol the host’s body. Upon sensing a broad variety of Pathogen-Associated or Danger-Associated Molecular Patterns (PAMPs and DAMPs, respectively), neutrophils extravasate from the blood and migrate to the “site of insult”. There, they deploy an impressive arsenal of antimicrobials to destroy invading pathogens. Neutrophils are very efficient phagocytes, engulfing and killing microorganisms by a combination of reactive oxygen species (ROS), proteases and other antimicrobial molecules (Amulic et al. 2012; Nordenfelt and Tapper 2011). They also release these molecules via degranulation, a process where the neutrophil’s storage granules fuse with the plasma membrane to release their content into the extracellular space (Lacy 2006). Another defense pathway of neutrophils is the release of Neutrophil Extracellular Traps (NETs), structures of DNA and proteins which neutrophils form upon different stimuli and via various pathways (Burgener and Schroder 2020; Brinkmann et al. 2004).

The antimicrobial functions of neutrophils are beneficial to the host, as illustrated by the finding that patients with deficiencies in neutrophil maturation or function suffer from recurrent bacterial and fungal infections (Klein 2011; Liew and Kubes 2019). However, as neutrophils arrive at inflammatory sites in considerable numbers and release a range of antimicrobials with a broad spectrum of action, they have the potential to damage host cells. As a result, neutrophil activation can also be associated with host pathology. Examples include neutrophil (over)activation in respiratory pathologies such as COPD or ARDS (Cowburn et al. 2008), in various cancers (Moses and Brandau 2016) and infectious diseases such as malaria (Knackstedt et al. 2019) or COVID-19 (Reusch et al. 2021; Schurink et al. 2020).

One difficulty working with neutrophils *ex vivo* is that the mature cells isolated from blood or from an inflammatory site are relatively short-lived, prone to activation and not amenable to genetic manipulation. This limits genetic experiments with human primary cells to studies on neutrophils from patients with known mutations. Human neutrophil cell lines are available (Rincón, Rocha-Gregg, and Collins 2018), however these do not recapitulate all of the functions of primary cells. One option to study genetically modified primary neutrophils is the use of mouse models. Indeed, there are neutrophil-specific Cre recombinase lines such as MRP8-Cre (Passegué, Wagner, and Weissman 2004) or Ly6G-Cre (Hasenberg et al. 2015) allowing conditional mutation of a gene of interest in the neutrophil population. These Cre lines, along with experiments where mouse neutrophils are depleted with anti-Ly6G-antibodies (Daley et al. 2008), have proven enormously valuable in understand the contribution of neutrophils to clearing infections, as well as in various pathologies *in vivo*.

While mouse neutrophils have been used extensively to study neutrophil function, it is not completely clear how well results from mouse studies translate to human neutrophils. In both species neutrophils are essential for defense against infections, but there are notable differences between mouse and human cells. One example is the proportion of neutrophils in circulation. Human neutrophils contribute to around 50-70% of circulating leukocytes while murine neutrophils make up around 10-25%, as discussed previously (Mestas and Hughes 2004; Doeing, Borowicz, and Crockett 2003). Another example is the expression of surface proteins, for example the bona fide marker for murine neutrophils (Ly6G) is absent on human cells (Lee et al. 2013).

We wanted to ask how the observed differences between mouse and human neutrophils may relate to actual functional differences. One confounding factor to consider is that most work with human cells has been done using neutrophils derived from blood. *Ex vivo* work with mouse cells, however, normally uses neutrophils from bone marrow or from the peritoneal cavity. Since neutrophils are generated in the bone marrow the population here may be less mature than those found in the blood. Conversely, neutrophil recruitment to the peritoneal cavity requires stimulation, such as casein or thioglycolate injection. Thus, peritoneal neutrophils may have a partially activated phenotype. It is therefore unclear whether differences observed between *ex vivo* mouse and human neutrophils are a true representation of species-specific differences or whether they reflect the different “states” of the cell or tissue location. For example, the release of neutrophils from bone marrow and transmigration to the inflammatory site is likely to be accompanied by activation. While studies have analysed the proteomes of neutrophils from human blood (Hoogendijk et al. 2019; Long et al. 2022; Grabowski et al. 2019; Rieckmann et al. 2017; Linder et al. 2023) and mice (Adrover et al. 2020), a systematic and quantitative comparison of neutrophils from different murine tissue sources and human blood is not available.

Here we used quantitative high-resolution proteomics to systematically understand how tissue location, infection or species impact the neutrophil protein landscape. We mapped three murine neutrophil populations; bone marrow, blood and peritoneal cavity, two of them commonly used for tissue culture experiments, revealing that the proteomes of these murine neutrophils are very similar, irrespective of the tissue source. Notable differences between murine populations were expression of some neutrophil transcription factors and an upregulation of chemokine and cytokine production as well as anti-apoptotic proteins in cells from the peritoneal cavity. Having mapped the core similarities and differences between mouse neutrophil populations we next examined the impact of a systemic fungal infection on the mouse bone marrow neutrophil proteome. We reveal that this infection triggers more than 1000 proteins to change in abundance. Lastly, we wanted to explore how similar mouse and neutrophil populations are by mapping the proteomes of neutrophils from blood. We reveal large differences between species including their total protein mass and some of the key neutrophil effector molecules. Together this data provides a valuable resource for comparing tissue-specific, infection-induced and species-specific neutrophil proteomes and provides new biological insight into model systems and their phenotypes. This comprehensive data set is freely available and easy to interrogate using the Immunological Proteome Resource (ImmPRes: immpres.co.uk).

## Results

### Profiling mouse bone marrow, blood and peritoneal neutrophil proteomes

Using quantitative mass spectrometry we characterised the proteomes of 3 key murine neutrophil populations: bone marrow, blood and peritoneal cavity. Bone marrow and blood neutrophils were taken from unstimulated mice, while peritoneal neutrophils were harvested following i.p. casein injection, a common method for inducing peritoneal neutrophil influx (Swamydas et al. 2015). Neutrophil populations were isolated to a purity >98% by fluorescent activated cell sorting (gating strategy shown in Supplementary Figure 1). We identified over 5800 proteins across the 3 populations (Fig 1A). The total cellular protein mass of bone marrow, blood and peritoneal neutrophils was approximately 40 pg/cell, with no major difference between populations (Fig 1B). Using the proteomics ruler (Wiśniewski et al. 2014) we estimated the copy number per cell for each protein, revealing a dynamic range of protein abundance from 100 copies per cell up to almost 100 million copies per cell (Supplementary File 1). Ranking all proteins by abundance revealed that 9 proteins account for 50% of all protein molecules detected within the cell and 54 proteins account for 75% of all molecules, in neutrophils isolated from bone marrow (Fig. 1C). This distribution looked similar across neutrophils from bone marrow, blood and peritoneal cavity (Supplementary Figure 2). Neutrophils store many of their effector molecules in granules. Proteins targeted to granules do not carry a specific signal peptide, instead they are loaded into granules during differentiation. The granule content reflects the transcriptional and translational program during the time of formation. Granules can roughly be divided into primary (or azurophilic), secondary (or specific), tertiary (or gelatinase) granules and secretory vesicles (Borregaard 2010), with ficolin granules forming a subset of secretory vesicles (Rørvig et al. 2009). Given the importance of granule proteins in neutrophil function we explored this group in more detail using an annotation of the granule subsets (Rørvig et al. 2013). Included in the most abundant neutrophil proteins were S100A8 and S100A9, which are annotated to secretory vesicles, and Neutrophil Granule Protein (NGP) which is also suggested to reside in murine neutrophil granules (Moscinski and Hill 1995). Together these 3 proteins accounted for almost 20% of all protein molecules expressed by the neutrophil (Fig 1C). We found that approximately 40% of the total protein molecules expressed by neutrophils could be assigned to granule subsets, with secretory vesicles accounting for the greatest proportion of the neutrophil protein content (Fig. 1D, E). This distribution was very similar between neutrophils isolated from different tissue sources (Fig. 1E). Next, we looked at proteins belonging to key cellular compartments such as the nuclear envelope, mitochondria and ribosome. We found no major difference between the protein content of the nuclear envelope or the mitochondria when comparing neutrophils from bone marrow, blood or peritoneal cavity (Fig. 1F). However, the ribosomal protein content was higher in bone marrow neutrophils compared with the other populations. The transition to mature neutrophils has been shown to correspond to reduced expression of proteins involved in mRNA translation (Hoogendijk et al. 2019). We confirm that when neutrophils leave the bone marrow and enter the blood stream, the proportion of their proteomes dedicated to ribosomal processes drops and remains lower in the peritoneal neutrophils (Fig. 1F).

**Figure 1.**
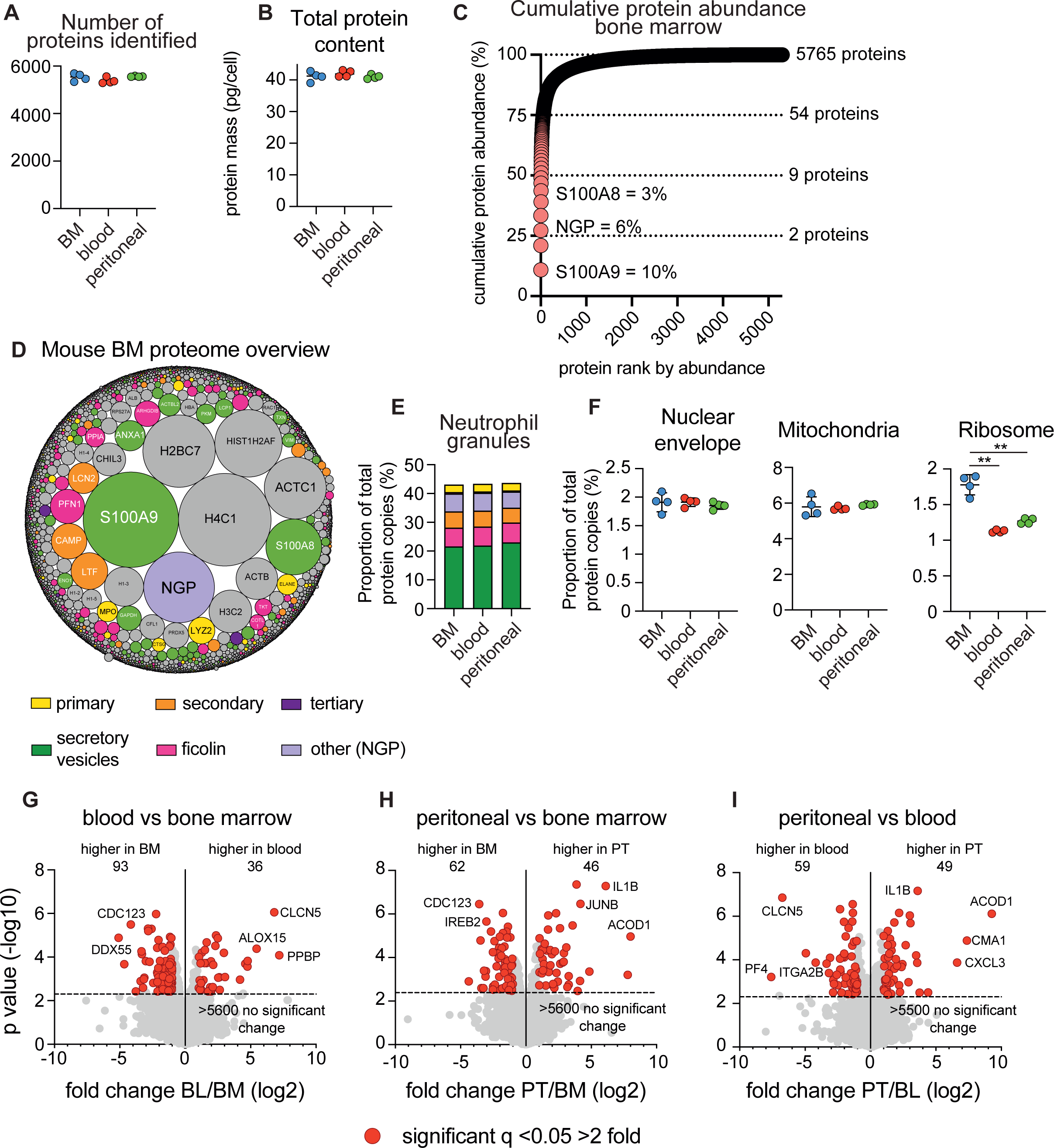
Mapping mouse tissue resident neutrophil proteomes. High-resolution quantitative mass spectrometry was used to characterise the proteomes of mouse neutrophils isolated from the bone marrow, blood and peritoneal cavity. (A) Number of proteins identified in each sample. (B) Total protein content of neutrophil populations. (C) Cumulative protein abundance for the bone marrow neutrophil proteome. Proteins were ranked according to their absolute copy number per cell. The number of proteins that make up 25%, 50%, 75% and 100% of the proteome is shown. (D) Overview of the mouse bone marrow proteome. Each protein is represented by a circle, with circle size indicating relative abundance. Granule proteins are subdivided into subsets (Rørvig et al. 2013) while non-granule proteins are coloured grey. (E) The contribution of granule protein subsets to the total protein molecules expressed by neutrophils. (F) The contribution of nuclear envelope proteins (Gene Ontology term 0005635), mitochondrial proteins (Gene Ontology term 0005739) and ribosomal proteins (Kyoto Encyclopaedia of Genes and Genomes annotation 03010) to the total protein molecules expressed by neutrophils. (G) Comparison of bone marrow and blood neutrophils (H) bone marrow and peritoneal neutrophils and (I) blood and peritoneal neutrophils. For the volcano plots proteins were deemed to be significantly changing with a q value <0.05 and a fold change >2 (highlighted in red). The horizontal dashed line on volcano plots indicates q = 0.05. BM: bone marrow, BL: blood, PT: peritoneal cavity. For each population 4 biological replicates were generated. Dot plots in a, b, and f show the mean +/- standard deviation. For 1F statistical significance was determined using an unpaired, unequal variance t-test with Welch’s correction. ** indicates p<0.01.

Next, we compared neutrophil proteomes to identify proteins that were differentially abundant between populations. The majority of the neutrophil proteome was unchanged between populations, with over 5500 proteins showing no significant difference in abundance between tissue locations, suggesting a high degree of proteome conservation between tissue resident neutrophils. However, we did identify proteins significantly different between the 3 populations. 129 proteins were differentially abundant between bone marrow and blood neutrophils, 108 proteins when comparing bone marrow and peritoneal cavity and 108 proteins when comparing blood and peritoneal cavity (Fig. 1G-I).

Enrichment analysis on differentially abundant proteins was used to identify core cellular processes that were significantly impacted by tissue location. Neutrophils isolated from the peritoneal cavity had elevated levels of proteins linked to the innate immune response including interleukins and chemokines. CXCL1, CXCL2, CXCL3, IL1B and IL36G were all found at higher levels in peritoneal neutrophils (Fig 2A). Neutrophils isolated from the bone marrow were enriched for proteins implicated in mRNA translation, and an analysis of ribosomal proteins showed elevated levels in bone marrow derived neutrophils (Fig 2B), explaining the higher ribosomal protein mass observed in Fig 1F. Neutrophils isolated from blood showed enrichment in proteins involved in cell adhesion and platelet aggregation including a number of fibrinogen molecules and the chemokine CCL6 (Fig 2C). The detection of platelet proteins could be the result of neutrophil-platelet aggregation (Reyes et al. 2021) or result from the uptake of platelet proteins by neutrophils. We also identified a selection of cell surface receptors that were differentially expressed depending on tissue location. Both components of the integrin complex ITGB3/ITGA2B were found at elevated levels in blood derived neutrophils (Fig 2D). This difference in integrin abundance was selective, and most integrin subunits were not differentially expressed between tissue locations. We also found the adhesion and homing receptor L-selectin at higher levels in blood neutrophils compared with bone marrow or peritoneal cavity (Fig 2D). Circulating neutrophils express high levels of L-selectin and it is shed in response to a range of stimuli and during transendothelial migration (Rahman et al. 2021; Ivetic 2018). Supporting this, we found over 12,000 copies per cell of L-selectin in blood neutrophils and this dropped to around 3,000 copies in peritoneal neutrophils (Fig 2D). Neutrophils isolated from the peritoneal cavity had increased levels of the C-type lectin-like domain containing receptors CLEC4D, 4E (Mincle) and 7A (Dectin-1) (Fig 2D).

**Figure 2.**
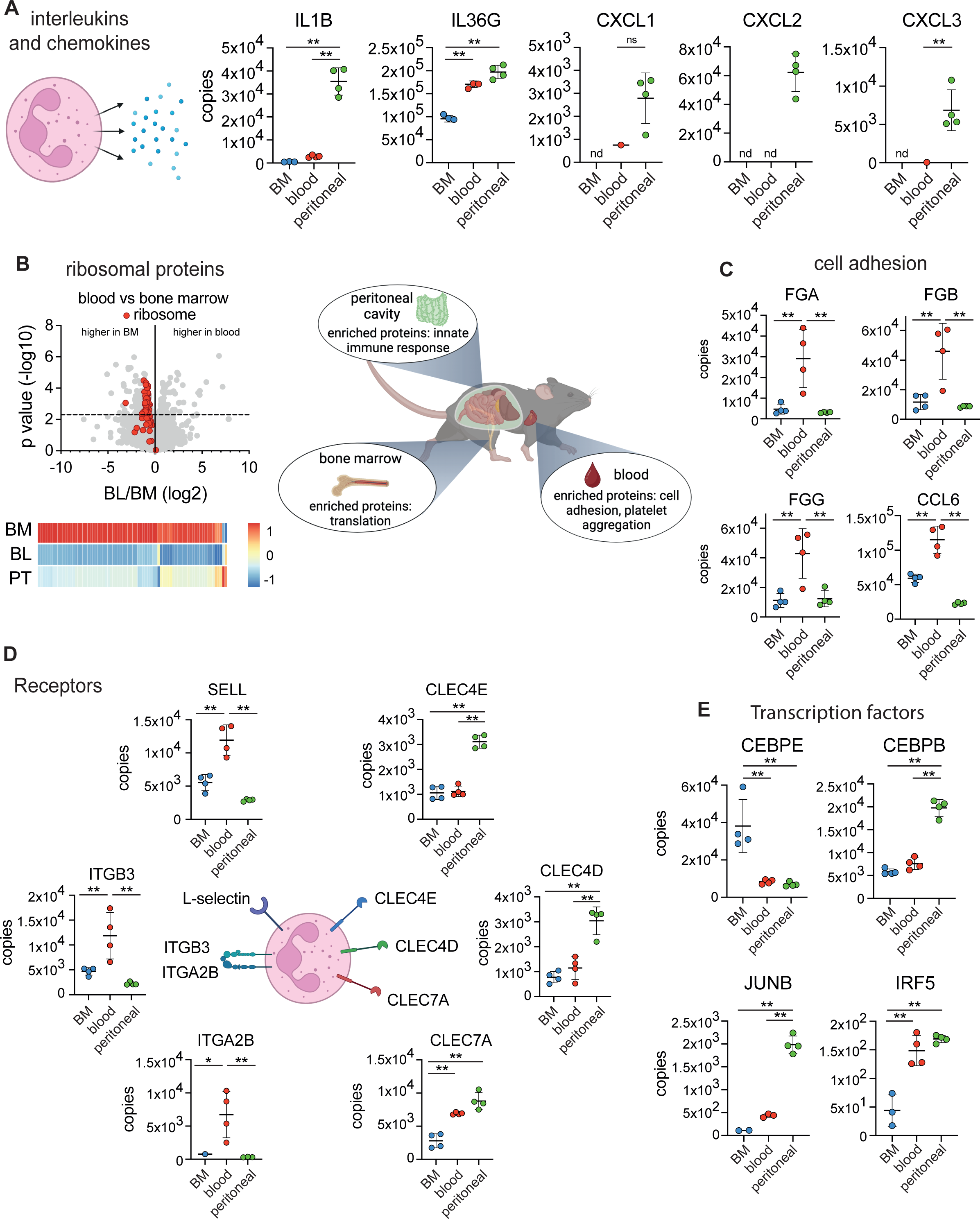
Tissue-specific neutrophil protein signatures. Neutrophils isolated from different tissues show enrichment in key processes. (A) The expression of innate immune response proteins C-X-C Motif Chemokine Ligand 1, 2 and 3 (CXCL1-3), Interleukin 1 Beta (IL1B) and Interleukin 36 Gamma (IL36G) in neutrophil populations. (B) The expression profile of ribosomal proteins (Kyoto Encyclopaedia of Genes and Genomes annotation 03010) in neutrophil populations. (C) The expression profile of a selection of proteins involved in cell adhesion in neutrophil populations. Fibrinogen Alpha Chain (FGA), Fibrinogen Beta Chain (FGB), Fibrinogen Gamma Chain (FGG) and Chemokine (C-C motif) ligand 6 (CCL6). (D) Protein copy numbers for a selection of cell surface receptors. L-selectin (SELL), C-Type Lectin Domain Family 4 Member E and D (CLEC4E and D), C-Type Lectin Domain Containing 7A (CLEC7A), Integrin Subunit Beta 3 (ITGB3) and Integrin Subunit Alpha 2b (ITGA2B). (E) The expression profile of 4 key transcription factors. CCAAT Enhancer Binding Protein Epsilon (CEBPE), CCAAT Enhancer Binding Protein Beta (CEBPB), JunB Proto-Oncogene AP-1 transcription factor subunit (JUNB) and Interferon Regulatory Factor 5 (IRF5). For each population 4 biological replicates were generated. Dot plots in a, c, d and e show the mean +/- standard deviation. BM: bone marrow, BL: blood, PT: peritoneal cavity. Statistical significance was determined using Limma with eBayes. The horizontal dashed line on volcano plot indicates q = 0.05. nd indicates when a protein was not detected, * indicates p<0.05, ** indicates p<0.01, ns indicates not significant.

Lastly, we analysed the profile of proteins with transcription factor or DNA binding activities. Over 150 proteins annotated as having transcription factor or DNA binding activity were identified within the neutrophil populations but there were few differences between tissue resident neutrophils (Supplementary Figure 3). However, a small number of transcription factors that are critical for neutrophil function were differentially expressed according to tissue. The C/EBP family of transcription factors are essential for neutrophil maturation as discussed in (Borregaard 2010). CEBPE has a role in the transcription of granule proteins and loss of expression of CEBPE leads to a condition called specific granule deficiency (Gombart et al. 2001). CEBPE was found at almost 40,000 copies per cell in bone marrow derived neutrophils but less than 10,000 copies per cell in blood and peritoneal derived neutrophils (Fig 2E). CEBPA is essential for homeostatic neutrophil maturation but can be replaced by CEBPB during emergency granulopoiesis (Hirai et al. 2006). While we didn’t detect CEBPA in our data set, we found that CEBPB was elevated in neutrophils isolated from the peritoneal cavity. CEBPB was found at approximately 6,500 copies in bone marrow and blood derived neutrophils but around 20,000 copies in peritoneal neutrophils (Fig 2E). JUNB was expressed most highly in peritoneal neutrophils while IRF5 showed greatest levels of expression in blood and peritoneal neutrophils (Fig 2E). Both JUNB and IRF5 are important for neutrophil effector function and regulate the expression of inflammatory molecules in neutrophils (Ericson et al. 2014; Ai and Udalova 2020; Khoyratty et al. 2021).

### The impact of systemic infection on mouse bone marrow neutrophils

Laboratory mice are commonly housed under specific pathogen-free conditions. Having shown that the bulk of the protein landscape is conserved between bone marrow, blood and peritoneal cavity resident neutrophils under these conditions, we decided to explore the impact of a systemic infection on murine bone marrow neutrophils. Mice were infected intravenously with *Candida albicans* (or PBS as a control) and neutrophils collected from bone marrow 5 days post infection, sorted by flow cytometry and analysed by mass spectrometry (Fig 3A, Supplementary Fig 4 and Supplementary File 1). Analysis of the total cellular protein mass revealed that neutrophils isolated from infected mice had a total protein mass similar to those from non-infected mice, 55 pg/cell for infected mice versus 45 pg/cell for non-infected (Fig 3B). Over 5,000 proteins showed no significant change in abundance in response to infection. However, we identified 1004 proteins that increased in abundance and 59 proteins that decreased in abundance in infected versus non-infected mice (Fig 3C). Those proteins found at increased abundance included proteins involved in translation, DNA replication and cell cycle. A closer look revealed that ribosomal and mitochondrial-ribosomal proteins were significantly increased in abundance and the proportion of total protein molecules dedicated to these compartments was elevated in response to infection (Fig 3D and 3E). A range of proteins involved in DNA replication/cell cycle were found at higher abundance in neutrophils from infected mice including components of the DNA replication fork complex such as the Minichromosome Maintenance Complex Components (MCM) enzymes (Fig 3F), DNA Ligase 1 and 3 (Supplementary Fig 5) and DNA Polymerase Alpha 1 and 2 (Supplementary Fig 5). It is likely that systemic infection induced emergency granulopoiesis, and an increase in immature neutrophils within the bone marrow. While Ly6G was used as a marker to collect mature neutrophils by cell sorting, the elevated levels of ribosomal and DNA replication proteins in bone marrow neutrophils from infected mice may be the result of the presence of some immature neutrophils within the sorted population. Molecules used for environmental sensing were also increased in abundance in neutrophils isolated from infected mice including Transferrin Receptor (TFRC), Integrin Subunit Alpha 4 (ITGA4) but not its partner Integrin Subunit Beta 1 (ITGB1), C-C Motif Chemokine Receptor 1 (CCR1), Sialic Acid Binding Ig Like Lectin 12 (SIGLEC12) and the intracellular pattern recognition receptor NLR Family CARD Domain Containing 4 (NLRC4) (Fig 3G). One notable group of molecules associated with neutrophils isolated from infected mice were the acute phase proteins. Serum amyloid proteins SAA1, SAA2 and APCS were only detected in neutrophils isolated from infected mice (Fig 3H). None of the major neutrophil effector molecules Elastase (ELANE), Proteinase 3 (PRTN3), Cathepsin G (CTSG) and Myeloperoxidase (MPO) changed significantly in response to infection (Fig 3I).

**Figure 3.**
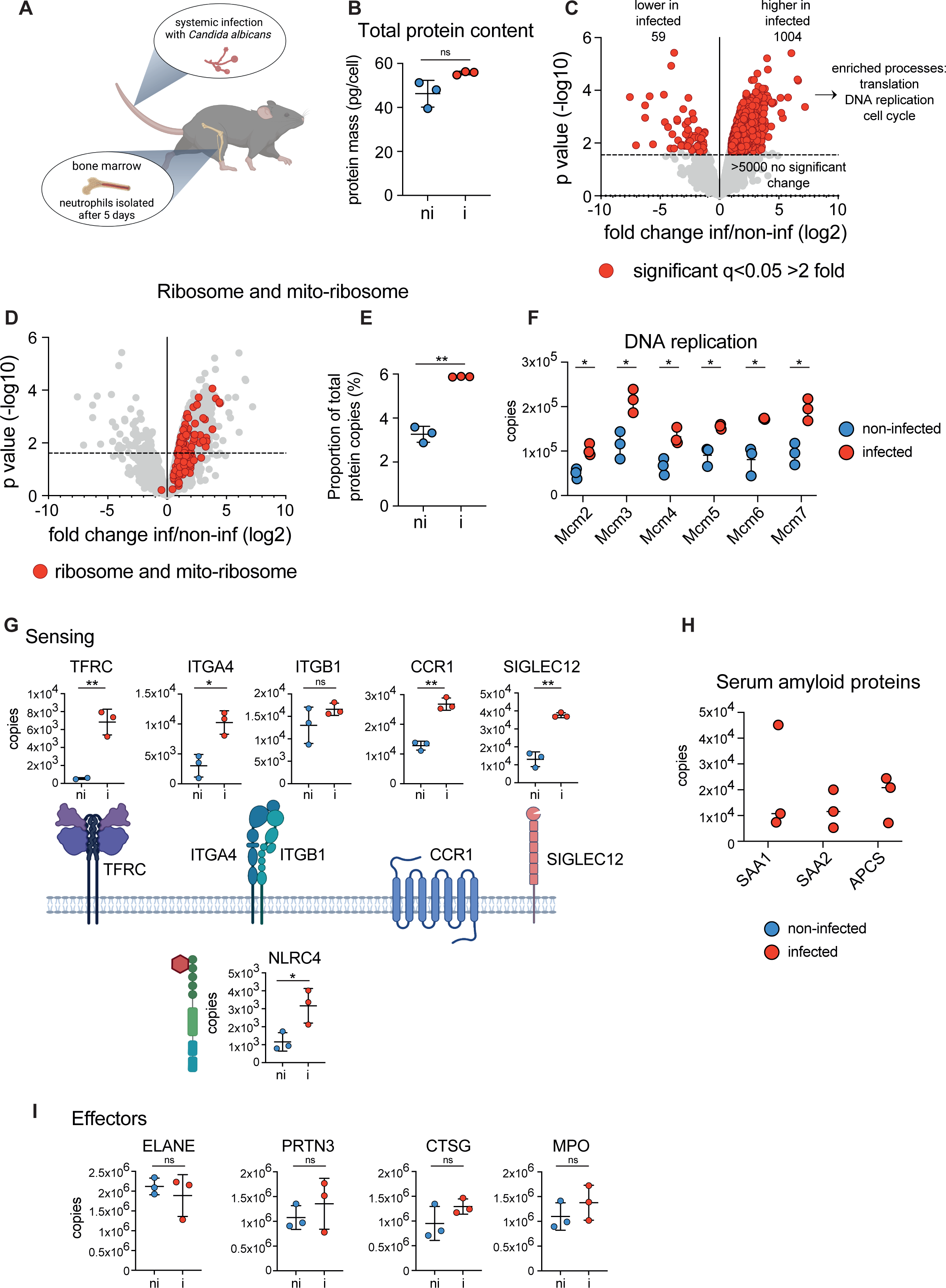
Systemic infection impacts core cellular processes and sensors in bone marrow neutrophils. (A) Mice were infected with *Candida albicans* and bone marrow neutrophils analysed by mass spectrometry 5 days post-infection. (B) Total protein content of neutrophil populations. (C) Comparison of bone marrow neutrophils isolated from infected versus non-infected mice. Proteins were deemed to be significantly changing with a q value <0.05 and a fold change >2 (highlighted in red). The horizontal dashed line on volcano plots indicates q = 0.05. (D and E) Expression profile of ribosomal and mitochondrial-ribosomal proteins in infected versus non-infected mice and the contribution that these proteins make to the total cellular protein molecules expressed. (F) The expression of MCM (Minichromosome Maintenance Complex Component) proteins in neutrophils isolated from the bone marrow of non-infected and infected mice. (G) The impact of *C. albicans* infection on the expression of sensing molecules in bone marrow neutrophils: Transferrin Receptor (TFRC), Integrin Subunit Alpha 4 (ITGA4), Integrin Subunit Beta 1 (ITGB1), C-C Motif Chemokine Receptor 1 (CCR1), Sialic Acid Binding Ig Like Lectin 12 (SIGLEC12) and NLR Family CARD Domain Containing 4 (NLRC4). (H) The abundance of serum amyloid proteins in neutrophils isolated from the bone marrow of non-infected and infected mice. (I) The impact of *C. albicans* infection on 4 key neutrophil effector molecules: Elastase (ELANE), Proteinase 3 (PRTN3), Cathepsin G (CTSG) and Myeloperoxidase (MPO). For each population 3 biological replicates were generated. Dot plots show the mean +/- standard deviation. For 3C, D, F, G and I statistical significance was determined using Limma with eBayes. ns indicates not significant while * indicates p<0.05, ** indicates p<0.01. For 3B and E statistical significance was determined using an unpaired, unequal variance t-test with Welch’s correction.

### Species-specific differences in blood derived neutrophils

Lastly, we wanted to explore the similarities and differences between murine and human neutrophils. Peripheral blood neutrophils are the most easily accessible source of human neutrophils and are commonly used for studying neutrophil function and disease phenotyping. We analysed human neutrophils derived from venous blood using the same mass spectrometry approach that was employed for mouse populations and directly compared human and mouse blood neutrophils. The average number of proteins identified was 4950 in human blood neutrophils, compared to 5384 proteins in mouse blood neutrophils (Fig 4A and Supplementary File 1). Interestingly, despite identifying slightly fewer proteins in the human neutrophils compared to mouse, we found a consistent increase in the total protein content of human neutrophils. Human blood neutrophils had a total protein mass over 60 pg/cell compared with 40 pg/cell for mouse blood neutrophils (Fig 4B). We analysed the profile of blood neutrophils by flow cytometry and found a considerable difference in their side scatter profile with human blood neutrophils showing a greater side scatter profile versus mouse neutrophils (Fig 4C and D). We explored which proteins were responsible for this difference in total cellular protein mass and found that human neutrophils contain significantly more granule proteins than mouse cells (Fig 4E and F). Primary/azurophilic granule proteins were particularly increased in human cells (Fig 4E and F). This was not limited to proteins known to be absent in mouse such as alpha-defensins or azurocidin (Hidalgo et al. 2019), but also included major neutrophil effector proteins stored in azurophilic granules, such as myeloperoxidase (MPO) and the serine proteases elastase (ELANE), proteinase 3 (PRTN3) and cathepsin G (CTSG) (Fig 4E and G). For example, CTSG was found at approximately 1.5 million copies per cell in mouse blood neutrophils but over 30 million copies per cell in human blood neutrophils. ELANE was found at approximately 3 million copies per cell in mouse blood neutrophils but over 20 million copies per cell in human blood neutrophils (Fig 4G). PRTN3, ELANE and CTSG are the major neutrophil effector proteases. However, not all proteases were found in greater abundance in human cells. For example, Cathepsin B and D (CTSB and CTSD) were more abundant in mouse blood neutrophils compared with human blood neutrophils (Fig 4G). Also, the apoptotic caspases Caspase-9 (CASP9) and Caspase-3 (CASP3) were found at higher levels in murine neutrophils (Fig 4G). Besides proteases, an examination of the NADPH oxidase components revealed less striking differences between mouse and human neutrophils. While a number of these proteins were significantly different between mouse and human, in most cases the differences were less striking, suggesting that not all effector proteins are equally affected (Fig 4H).

**Figure 4.**
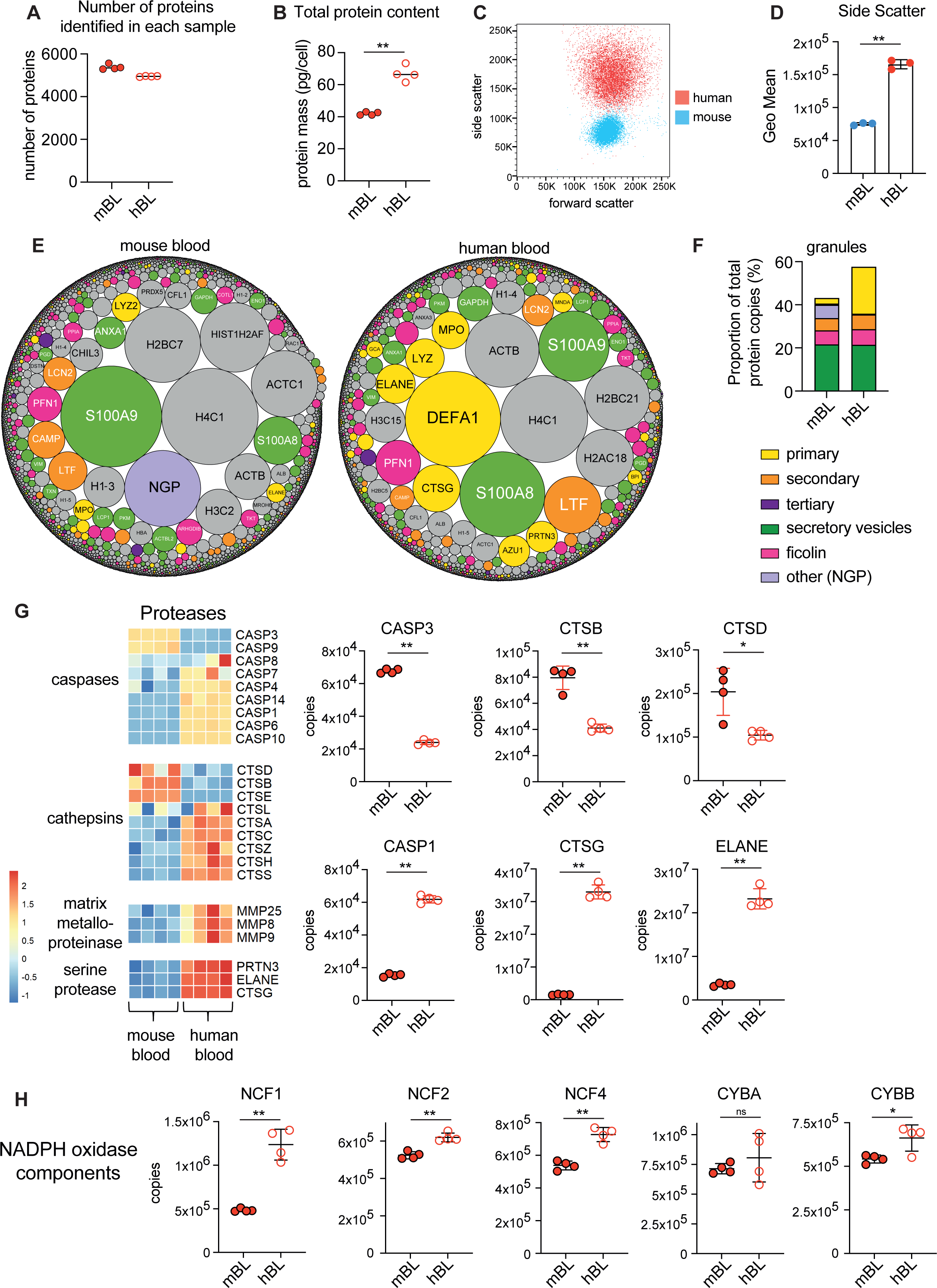
Species-specific neutrophil protein signatures. Comparison of mouse and human blood neutrophils. (A) Number of proteins identified in each sample. (B) Total protein content of neutrophil populations. (C) Analysis of forward and side scatter profile of human and mouse blood neutrophils by flow cytometry. (D) Geometric mean of neutrophil side scatter for human and mouse cells. (E) Overview of mouse and human blood neutrophil proteomes. Each protein is represented by a circle, with circle size indicating relative abundance. Granule proteins are subdivided into subsets (Rørvig et al. 2013) while non-granule proteins are coloured grey. (F) The contribution of granule protein subsets to the total protein molecules expressed by mouse and human blood neutrophils. (G) The expression profile of a selection of proteases in mouse and human neutrophils. Caspase 1 (CASP1) Caspase 3 (CASP3), Cathepsin B (CTSB), Cathepsin G (CTSG), Cathepsin D (CTSD) and Elastase (ELANE). (H) The abundance of NADPH oxidase components in mouse and human neutrophils. For each population analysed by proteomics 4 biological replicates were generated. Dot plots show the mean +/- standard deviation. mBL: mouse blood, hBL: human blood. For flow cytometry analysis of human and mouse blood neutrophils 3 biological replicates were analysed. Statistical significance was determined using an unpaired, unequal variance t-test with Welch’s correction. * indicates p<0.05, ** indicates p<0.01.

Lastly, we looked at the abundance of interleukins and chemokines. We detected IL18 only in human blood neutrophils while IL1B was only detected in mouse blood neutrophils. IL16 was found in greater abundance in mouse blood neutrophils (Fig 5A). CXCL1, 2 and 6 were either increased in abundance or exclusively found in human neutrophils while CCL6 was only found in mouse neutrophils (Fig 5B), suggesting that human and mouse neutrophils differentially produce chemokines and cytokines which may impact their recruitment to inflammatory sites. Together our data reveals major differences between human and murine neutrophils, the most striking difference being a significant increase in primary granule protein content in human cells.

**Figure 5.**
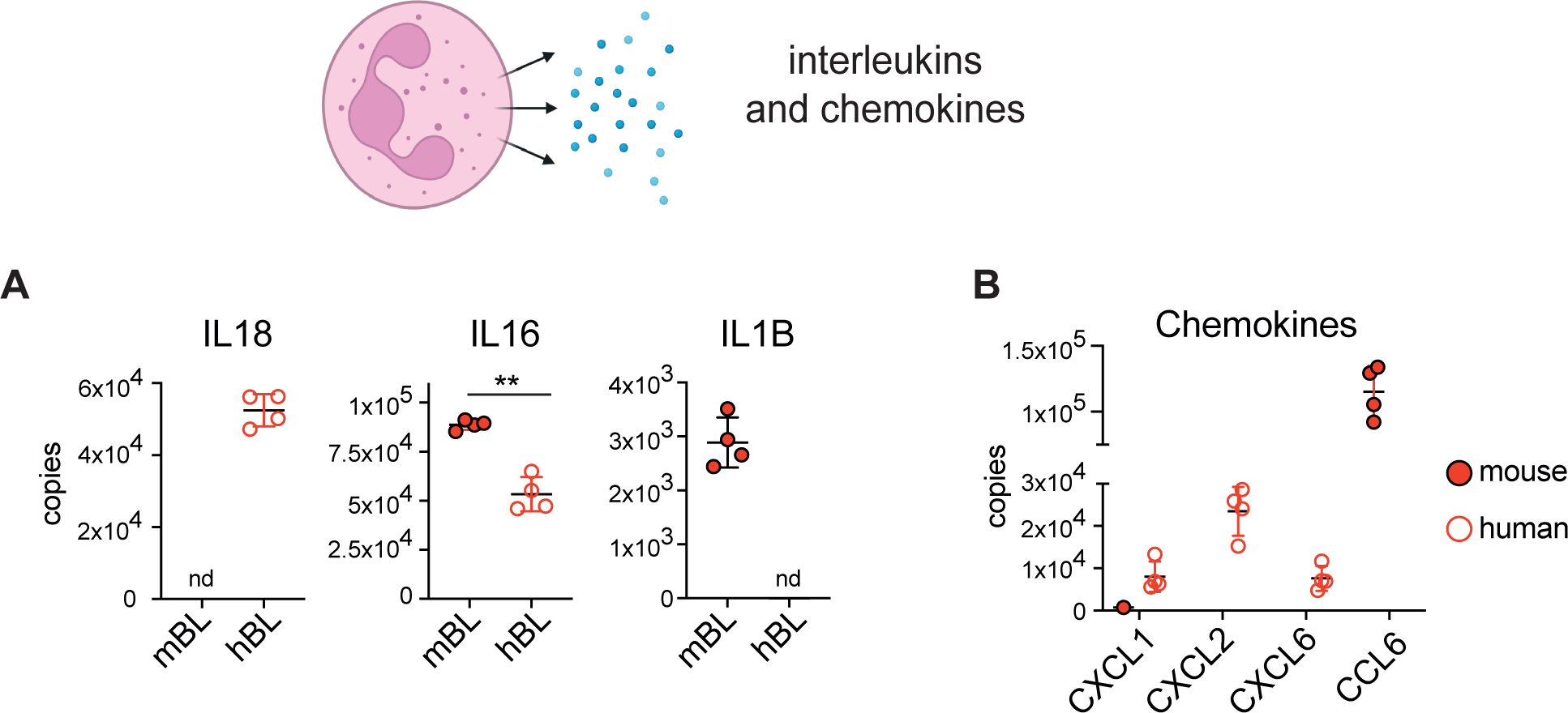
Human and mouse neutrophils differ in their expression of secreted molecules. (A) Interleukins and (B) chemokines that are differentially expressed in across species. Interleukin 18 (IL18), Interleukin 16 (IL16), C-X-C Motif Chemokine Ligand 1, 2, 6 (CXCL1,2 and 6) and Chemokine (C-C motif) ligand 6 (CCL6). For each population 4 biological replicates were generated. Dot plots show the mean +/- standard deviation. mBL: mouse blood, hBL: human blood. nd indicates when a protein was not detected. Statistical significance was determined using an unpaired, unequal variance t-test with Welch’s correction. * indicates p<0.05, ** indicates p<0.01.

## Discussion

This study provides a comprehensive resource for understanding the impact of tissue location, systemic infection and species on the neutrophil protein landscape. Using high sensitivity mass spectrometry we identified over 5000 proteins and estimated the copy number per cell for each protein across neutrophil populations. This approach allowed us to calculate the proportion of the neutrophil proteome dedicated to key processes. We provide novel insights into the core machinery, granule content, effector molecules and environmental sensors expressed by neutrophils.

Our first aim was to understand whether neutrophils from distinct tissue environments show unique protein signatures. We revealed that murine bone marrow, blood and peritoneal neutrophils are highly conserved in their core protein landscape with over 5000 proteins showing no difference in response to tissue niche. However, we identified 129 proteins that were differentially expressed when comparing bone marrow and blood neutrophils, 108 proteins when comparing bone marrow and peritoneal cavity and 108 proteins when comparing blood and peritoneal cavity. Bone marrow neutrophils showed elevated levels of proteins involved in translation, with ribosomal protein mass increased. Mature neutrophils are thought to possess all their effector proteins and to have relatively low translational activity. As the bone marrow is the site of neutrophil differentiation, the higher ribosomal content relative to blood and peritoneal neutrophils might reflect this transition to mature cells. Blood neutrophils were enriched in proteins involved in cell adhesion and platelet aggregation, which could either be through their association directly with platelets or by the direct uptake of platelet proteins by neutrophils. Blood neutrophils also showed elevated expression of the homing receptor L-selectin and integrin alpha-IIb/beta-3 (ITGA2B:ITGB3). Peritoneal neutrophils showed elevated levels of molecules associated with innate immune cell activation or immunity including interleukins and chemokines and also the pattern recognition receptors CLEC4D, CLEC4E and CLEC7A. (CLEC7A (Dectin-1) recognizes □-glucans and is involved in antifungal defense (Goodridge, Wolf, and Underhill 2009). CLEC4E (Mincle) binds glycolipids and has also been shown to be involved in defense against fungal species (Wells et al. 2008; Yamasaki et al. 2009), but can also recognize bacterial products such as Trehalose 6,6′-dimycolate from mycobacteria (Lee et al. 2012). CLEC4D and CLEC4E co-regulated in myeloid cells in response to immune challenge (Kerscher et al. 2016). It is likely that upregulation of these receptors in peritoneal neutrophils facilitates antimicrobial action of the cells at an inflammatory site.

When we explored the impact of systemic infection on the mouse bone marrow proteome, we noticed that the bulk of the neutrophil proteome remained unchanged. However, ribosomal proteins along with proteins involved in cell cycle and DNA replication were elevated in infected mice, perhaps through the production of considerable numbers of neutrophils upon infection. While, based on our sorting criteria, cells from non-infected and infected mice had the same levels of Ly-6G expression, it is possible that bone marrow neutrophils from infected mice are more immature, which could explain the above-mentioned findings. Interestingly, a number of serum amyloid A (SAA) proteins were found only in neutrophil samples from infected mice. SAA are known to increase considerably upon conditions of infection and inflammation and are predominantly produced by hepatocytes. SAA proteins were also found at elevated levels in the proteomes of neutrophils in response to COVID-19 (Long et al. 2022) (http://immpres.co.uk/). Neutrophils have previously been shown to scavenge extracellular proteins to fuel metabolism (Watts et al. 2021). However, it remains to be determined whether the detection of SAA proteins in neutrophils from infected mice in this study is the result of protein scavenging.

We also compared mouse and human blood neutrophils to identify species-specific differences. We observed a striking difference in the total protein mass of mouse and human neutrophils. The total protein mass of human cells was over 50% greater than mouse cells. Flow cytometry analysis revealed that human neutrophils have a greater side scatter profile while proteomics revealed a significantly higher abundance of primary granule proteins in human cells. As far as we are aware this is the first report of this difference between mouse and human neutrophils. Given that mice and humans follow different circadian rhythms and neutrophils spontaneously degranulate as they age (Adrover et al. 2020), one could hypothesize that the difference in granule content is the result of cells aging at different times. However, we suspect that this is not the case since the difference in granule content was predominantly seen in primary granules which are usually the last granules to be released. In contrast, mouse and human neutrophils devoted the same proportion of their protein mass to secretory vesicles, and these are among the first granules to release. Further studies are required to establish why human neutrophils have a higher primary granule content, but one hypothesis could be a requirement for human cells to have a greater cytotoxic capability due to the considerably larger tissue area that they are required to protect.

One limitation of our study is that we used bulk populations of neutrophils. Neutrophils are however not a homogeneous population and in future it may be possible to distinguish neutrophil subsets using specific markers, such as OLFM4 (Clemmensen et al., 2012). As mass spectrometry approaches advance and require less input material, future studies may address neutrophil heterogeneity at the protein level in more detail.

In summary, we believe that this work will help inform future neutrophil studies and will be a reference for understanding neutrophil functions and phenotypes. Our data is freely available and easy to interrogate using the Immunological Proteome Resource (immpres.co.uk), allowing users to compare protein abundance between model systems and across tissue locations.

## Methods

### Mice

For experiments comparing tissue resident neutrophils male wildtype C57BL/6 mice (Charles Rivers) aged between 8 and 14 weeks were used. For experiments examining the impact of *Candida albicans* infection male and female mice were used, with an equal gender balance across the experiment. Animals were kept in individually ventilated cages in a pathogen free facility at 21°C between 45 and 65% humidity and on a 12/12h light/dark cycle. Mice were provided with free access to food (R&M3, Special diet Services) and water. For isolation of peritoneal and bone marrow neutrophils mice were sacrificed using a rising concentration of CO_2_ and death subsequently confirmed by either cervical dislocation or exsanguination. For the isolation of blood neutrophils mice were anaesthetised using pentabarbitone (Euthatal) at 40mg/ml and dosed at 0.1ml per 10g body weight i.p. Blood was extracted by exsanguination via cardiac puncture whilst the animal was anaesthetised. For *Candida albicans* infection experiments mice were injected intravenously with either 1×10^5^CFU of *Candida albicans* (strain SC5314) or PBS as a control. Mice were monitored daily and culled after 5 days by CO_2_ as described above. All mice were maintained in the Biological Resource Unit at the University of Dundee using procedures approved by the University of Dundee Ethical Review Committee and under the authorization of the UK Home Office Animals (Scientific Procedures) Act 1986.

### Ethical approval for human blood work

This study was approved by the local ethics committee of the University of Dundee. All participants gave written informed consent.

### Isolation of pure populations of neutrophils

#### Mouse bone marrow neutrophils

Bone marrow was flushed from the tibias and femurs of mice in PBS under sterile conditions. Neutrophils were enriched from bone marrow using a negative selection immuno-magnetic isolation kit according to the manufacture’s protocol (Miltenyi Biotec). Enriched neutrophils were suspended in PBS with 1% FBS and incubated with mouse FC block at a concentration of 1 µg/million cells for 10 minutes on ice. Cells were then stained with Ly-6G FITC, CD11b APC and DAPI and a pure population of neutrophils (Ly6G and CD11b high) were isolated by cell sorting. Sorted cells were washed twice with PBS and cell pellets flash frozen in liquid nitrogen and stored at -80 °C until processing for mass spectrometry. For *C. albicans* infection study neutrophils were isolated from the bone marrow of infected and non-infected mice as described above but using the following markers: Ly-6G FITC, CD11b PE-Cy7 and DAPI.

#### Mouse blood neutrophils

Mouse blood (collected as described above) was collected into tubes containing EDTA as an anticoagulant. For each biological replicate blood was collected from 5 wildtype mice and pooled. The collected blood was topped up to 10ml with ice-cold PBS. To the Blood-PBS solution 40 ml of ice-cold red blood cell lysis buffer was added for a final concentration of 150mM ammonium chloride, 10mM sodium bicarbonate and 1.2mM EDTA. The sample was incubated on ice for 15 minutes and gently inverted every 5 minutes. After incubation the blood suspension was centrifuged at 300g for 10 minutes and washed once with PBS containing 2 % FBS and 1mM EDTA, before repeating the centrifugation step above. The cell pellet was suspended in PBS with 1% FBS and FC blocked, stained and sorted as described above.

#### Mouse peritoneal neutrophils

Casein was slowly dissolved in PBS at 80 to 90°C to give a 9% solution. It was then autoclaved and stored frozen at -80°C. To recruit neutrophils to the peritoneum, casein was injected intraperitoneally at a dose of 20 ml/kg. After 16 h mice were given a 2^nd^ i.p. injection of casein at the same dose and then sacrificed 3h later. Cells were collected from the peritoneum by injecting 3 ml of sterile PBS containing 0.02% (w/v) EDTA and then collecting the fluid from the peritoneal cavity. Cells were then pelleted by centrifugation for 5min and then resuspended in PBS with 1% FBS and FC blocked, stained and sorted as described above.

#### Human peripheral blood neutrophils

10 ml of blood was taken from 4 donors and collected into individual tubes containing EDTA as the anticoagulant. Neutrophils were isolated by immuno-magnetic negative selection using the EasySep Direct human neutrophil isolation kit (StemCell Technologies). A highly pure population of neutrophils was then generated by cell sorting on the basis of size, granularity, and autofluorescence based on forward/side scatter and autofluorescence as previously described (Dorward et al. 2013).

### Proteomics sample preparation

Cell pellets were lysed in 100 µl lysis buffer (5% SDS, 10 mM TCEP, 50 mM TEAB) and shaken at room temperature for 5 minutes at 1000 rpm, followed by boiling at 95 °C for 5 minutes at 500 rpm. Samples were then shaken again at RT for 5 minutes at 1000 rpm before being sonicated for 15 cycles of 30 seconds on/ 30 seconds off with a BioRuptor (Diagenode). Benzonase was added to each sample and incubated at 37 °C for 15 minutes to digest DNA. Samples were then alkylated with the addition of iodoacetamide to a final concentration of 20 mM and incubated for 1 hour in the dark at 22 °C. Protein concentration was determined using EZQ protein quantitation kit (Invitrogen) as per manufacturer instructions. For experiments comparing tissue resident neutrophils and human blood neutrophils protein clean-up and digestion was performed using S-TRAP micro columns following the manufacturers instructions (Protifi). Proteins were digested with trypsin at 1:10 ratio (enzyme:protein) for 2 hours at 47 °C. Digested peptides were eluted from S-TRAP columns using 50 mM ammonium bicarbonate, followed by 0.2 % aqueous formic acid and with a final elution using 50% aqueous acetonitrile. Eluted peptides were dried overnight before being resuspended in 40 µl 1 % formic acid ready for analysis by data independent acquisition mass spectrometry.

For experiments examining the impact of *Candida albicans* infection protein clean-up and digestion was performed using the SP3 method as described by (Hughes et al. 2014; Damasio et al. 2021). Peptides were fractionated by high pH reverse phase fractionation as described previously (Damasio et al. 2021). Samples were loaded onto a XbridgeTM BEH130 C18 column with 3.5 μm particles (Waters). Using a Dionex BioRS system, the samples were separated using a 25-min multistep gradient of solvents A (10 mM formate at pH 9 in 2% acetonitrile) and B (10 mM ammonium formate at pH 9 in 80% acetonitrile) at a flow rate of 0.3 ml min−1. Peptides were separated into 16 fractions and subsequently consolidated into 8 fractions. The fractions were dried, and peptides dissolved in 1% formic acid and analysed by liquid chromatography mass spectrometry.

### Mass spectrometry

For experiments comparing tissue resident neutrophils and human blood neutrophils, peptides were analysed by data independent acquisition (DIA), 1.5 µg of peptide from each sample was analysed. Peptides were injected onto a nanoscale C18 reverse-phase chromatography system (UltiMate 3000 RSLC nano, Thermo Scientific) and electrosprayed into an Orbitrap Exploris 480 Mass Spectrometer (Thermo Fisher). For liquid chromatography the following buffers were used: buffer A (0.1% formic acid in Milli-Q water (v/v)) and buffer B (80% acetonitrile and 0.1% formic acid in Milli-Q water (v/v). Samples were loaded at 10 μL/min onto a trap column (100 μm × 2 cm, PepMap nanoViper C18 column, 5 μm, 100 Å, Thermo Scientific) equilibrated in 0.1% trifluoroacetic acid (TFA). The trap column was washed for 3 min at the same flow rate with 0.1% TFA then switched in-line with a Thermo Scientific, resolving C18 column (75 μm × 50 cm, PepMap RSLC C18 column, 2 μm, 100 Å). Peptides were eluted from the column at a constant flow rate of 300 nl/min with a linear gradient from 3% buffer B to 6% buffer B in 5 min, then from 6% buffer B to 35% buffer B in 115 min, and finally to 80% buffer B within 7 min. The column was then washed with 80% buffer B for 4 min and re-equilibrated in 3% buffer B for 15 min. Two blanks were run between each sample to reduce carry-over. The column was kept at a constant temperature of 50oC.

The data was acquired using an easy spray source operated in positive mode with spray voltage at 2.445 kV, and the ion transfer tube temperature at 250oC. The MS was operated in DIA mode. A scan cycle comprised a full MS scan (m/z range from 350-1650), with RF lens at 40%, AGC target set to custom, normalised AGC target at 300%, maximum injection time mode set to custom, maximum injection time at 20 ms, microscan set to 1 and source fragmentation disabled. MS survey scan was followed by MS/MS DIA scan events using the following parameters: multiplex ions set to false, collision energy mode set to stepped, collision energy type set to normalized, HCD collision energies set to 25.5, 27 and 30%, orbitrap resolution 30000, first mass 200, RF lens 40%, AGC target set to custom, normalized AGC target 3000%, microscan set to 1 and maximum injection time 55 ms. Data for both MS scan and MS/MS DIA scan events were acquired in profile mode.

For experiments examining the impact of *Candida albicans*, peptides were analysed by data dependent acquisition (DDA). 1 µg of peptide from each fraction was analysed using a LTQ Orbitrap Velos (Thermo Fisher Scientific) as described in detail previously (Marchingo et al. 2020).

### Mass spectrometry data analysis

Raw DIA mass spec data files were searched using Spectronaut version 16.0.220606.53000. Data was analysed using a hybrid library approach. For mouse data, the hybrid library was created using two data sets: in-depth bone marrow fractionated neutrophil DDA proteomics data along with single shot DIA data for experimental samples (blood, bone marrow and peritoneal neutrophils). To generate an in-depth bone marrow neutrophil library, peptides from bone marrow neutrophils (isolated and prepared as described above) were fractionated by high pH reverse phase fractionation into 8 fractions and analysed by data dependent acquisition (as described for the *C. albicans* infection experiment above). Raw mass spec data files were searched using the Pulsar tool within Spectronaut using the following settings: 0.01 FDR at the protein and peptide level with digest rule set to ‘Trypsin’P’. A maximum of two missed cleavages and minimum peptide length of 7 amino acids was selected. Carbamidomethyl of cysteine was selected as a fixed modification while protein n-terminal acetylation and methionine oxidation were selected as variable modifications. The data was searched against a mouse database from the Uniprot release 2020 06. The database was generated using all manually annotated mouse SwissProt entries, combined with mouse TrEMBL entries with protein level evidence available and a manually annotated homologue within the human SwissProt database. To generate a library from experimental samples, blood, bone marrow and peritoneal neutrophil samples generated by DIA (described in the mass spectrometry section above) were analysed using the Pulsar tool within Spectronaut and using the same settings as were used for the fractionated bone marrow data above. Experimental DIA samples were then searched against both the fractionated DDA library and the experimental DIA library using the following identification settings: protein and precursor q-value set to 0.01 with an ‘Inverse’ decoy method and ‘Dynamic’ decoy limit strategy. The following quantification settings were used: ‘Quant 2.0’, the MS-Level quantity was set to ‘MS2’, imputation was disabled, major group Top N and minor group Top N were set as ‘False’ and cross run normalisation was set as ‘False’.

For human DIA data a hybrid library approach was used. The human hybrid library was created using two data sets: in-depth fractionated human blood neutrophil DDA proteomics data along with single shot DIA data for human blood experimental samples. To generate an in-depth human blood neutrophil library, peptides from human blood neutrophils (isolated and prepared by the s-trap method described above) were fractionated by high pH reverse phase fractionation into 16 fractions and analysed by data dependent acquisition. Raw mass spec data files were searched using the Pulsar tool within Spectronaut using the following settings: 0.01 FDR at the protein and peptide level with digest rule set to ‘Trypsin’P’. A maximum of two missed cleavages and minimum peptide length of 7 amino acids was selected. Carbamidomethyl of cysteine was selected as a fixed modification while protein n-terminal acetylation and methionine oxidation were selected as variable modifications. The data was searched against a human SwissProt database release 2020 07. To generate a library from experimental samples, human blood neutrophil data generated by DIA (described in the mass spectrometry section above) were analysed using the Pulsar tool within Spectronaut and using the same settings as were used for the fractionated blood data above. Experimental DIA samples were then searched against both the fractionated DDA library and the experimental DIA library using the following identification settings: protein and precursor q-value set to 0.01 with an ‘Inverse’ decoy method and ‘Dynamic’ decoy limit strategy. The following quantification settings were used: ‘Quant 2.0’, the MS-Level quantity was set to ‘MS2’, imputation was disabled, major group Top N and minor group Top N were set as ‘False’ and cross run normalisation was set as ‘False’.

For neutrophils from the *C. albicans* infection experiment raw mass spectrometry DDA data files were analysed using MaxQuant (Cox and Mann 2008) version 1.6.10.43. Proteins and peptides were identified using a hybrid database from the Uniprot release 2019 08. The hybrid database was generated using all manually annotated mouse SwissProt entries, combined with mouse TrEMBL entries with protein level evidence available and a manually annotated homologue within the human SwissProt database. The following search parameters were used: trypsin and LysC were selected as digestion enzymes with up to 2 missed cleavages; the false discovery rate was set at 1% for protein and peptide and the match between runs function was disabled. Methionine oxidation, protein N-terminal acetylation, deamidation (NQ) and Gln to pyro-Glu were selected as variable modifications while carbamidomethylation of cysteine was selected as a fixed modification. Proteins were removed from the data set which were categorised as ‘reverse’, ‘contaminant’ or ‘only identified by site’.

To estimate protein copy numbers per cell we used the proteomics ruler (Wiśniewski et al. 2014). Protein differential expression analysis was performed using R (v. 4.0.3) and p-values and fold changes were calculated using the Bioconductor package Limma (v 3.46.0) (Ritchie et al. 2015). The q-value was calculated with the Bioconductor package qvalue (v 2.22.0). Heatmaps were generated using the pheatmap package in R. Missing values were replaced with a value of 1 and all values were log transformed. Z scores were calculated as (x-mean(x))/sd(x) for each row of the dataset and heatmaps were visualized with cluster_cols set as false.

### Analysis of mouse and human blood neutrophils by flow cytometry

Mouse and human blood were extracted as described above and collected into anticoagulant tubes containing EDTA. Red blood cells (RBC) were lysed using a 1× RBC lysis buffer (eBioscience) according to the manufacturers instructions. Following RBC lysis the cell suspension was centrifuged at 300g at 4°C for 5 mins and then suspended in PBS with 1% FBS and FC blocked (as described above) and stained with CD66b FITC and DAPI for human cells and Ly6G FITC, CD11b APC and DAPI for mouse cells. Forward and side scatter profiles were analysed by flow cytometry.

## Data availability

Raw mass spec data files and result files are available from the ProteomeXchange data repository (http://proteomecentral.proteomexchange.org/cgi/GetDataset) and can be accessed with the following details: accession code PXD044568 for mouse bone marrow, blood and peritoneal cavity neutrophil populations, PXD044556 for mouse *C. albicans* infection neutrophils, PXD044569 for human blood neutrophils. All proteomics data is freely available on the Immunological Proteome Resource (ImmPRes: immpres.co.uk).

## Acknowledgments

The authors would like to thank Rosie Clarke and Arlene Rennie from the flow cytometry facility at the University of Dundee for cell sorting. We thank the biological sciences research unit and Sarah Thomson at the University of Dundee for support with animal work. We would also like to thank Borko Amulic (University of Bristol) and Arturo Zychlinsky (Max Planck Institute for Infection Biology Berlin) for providing feedback on the manuscript. Graphics used in figures were created using BioRender.com. This research was supported by an AMS Springboard Award (SBF006_1137) to GS. AH was supported by a Wellcome Trust Principal Research Fellowship, awarded to Doreen Cantrell (205023/Z/16/Z).

## Figure Legends

**Supplementary Figure 1.**
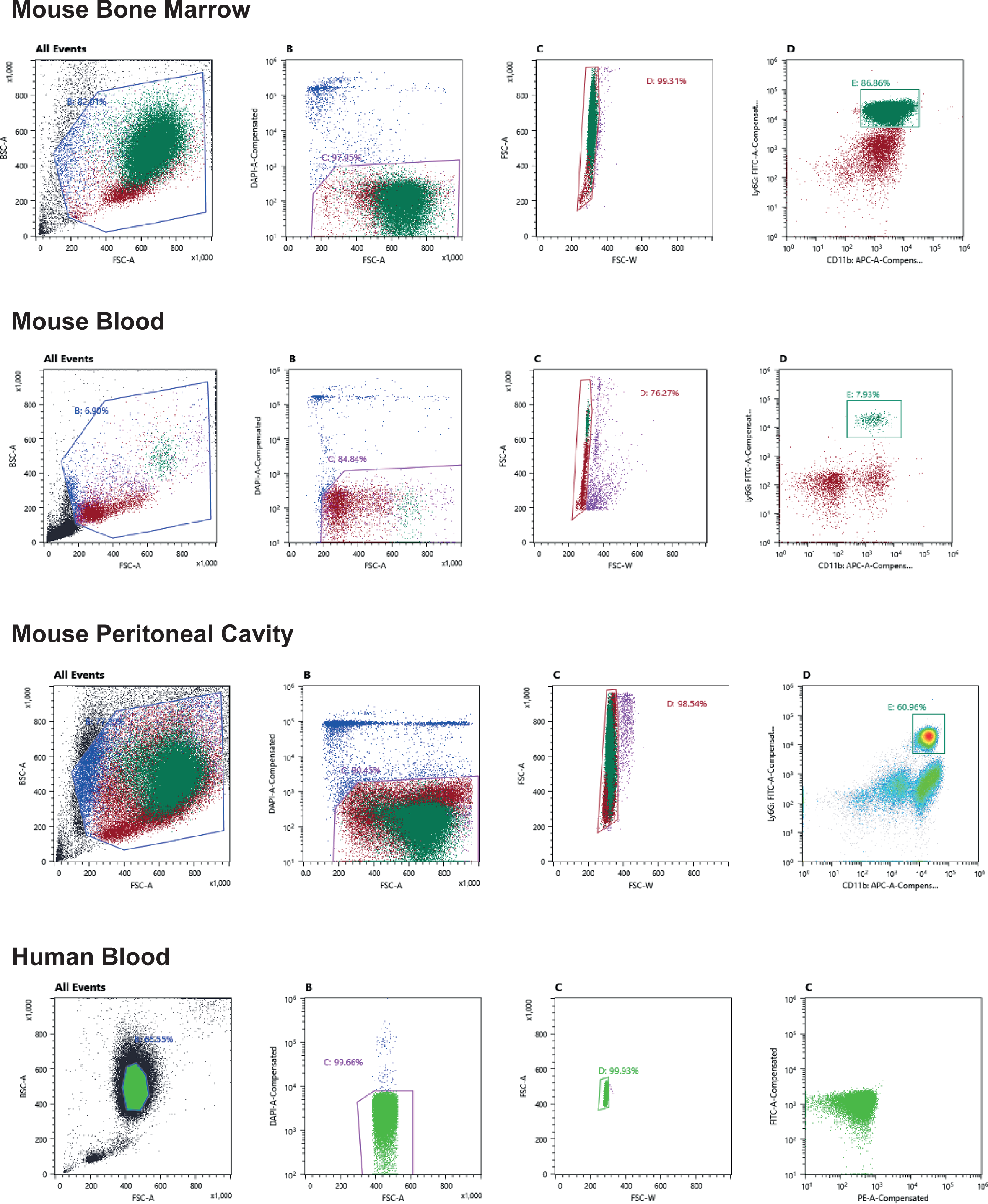
Gating strategy for fluorescent activated cell sorting pure neutrophil populations from mouse bone marrow, blood and peritoneal cavity and from human blood. Mouse neutrophil populations were stained with Ly-6G FITC, CD11b APC and DAPI and a pure population of neutrophils (Ly6G and CD11b high) were isolated by cell sorting. Human blood neutrophils were sorted on the basis of size, granularity, and autofluorescence based on forward/side scatter and autofluorescence as previously described (Dorward et al. 2013).

**Supplementary Figure 2.**
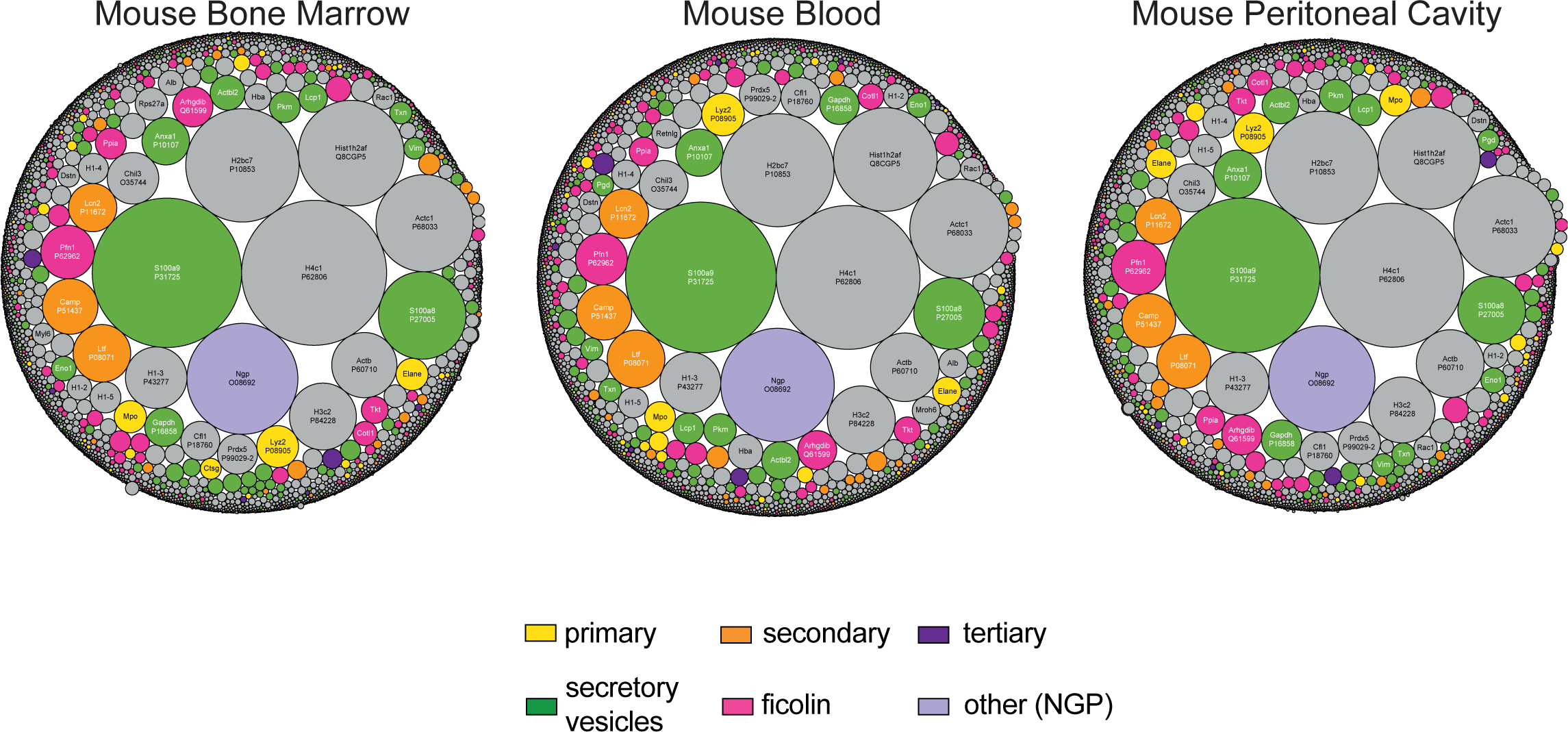
Overview of mouse bone marrow, blood and peritoneal cavity neutrophil proteomes. Each protein is represented by a circle, with circle size indicating relative abundance. Granule proteins are subdivided into subsets (Rørvig et al. 2013) while non-granule proteins are coloured grey.

**Supplementary Figure 3.**
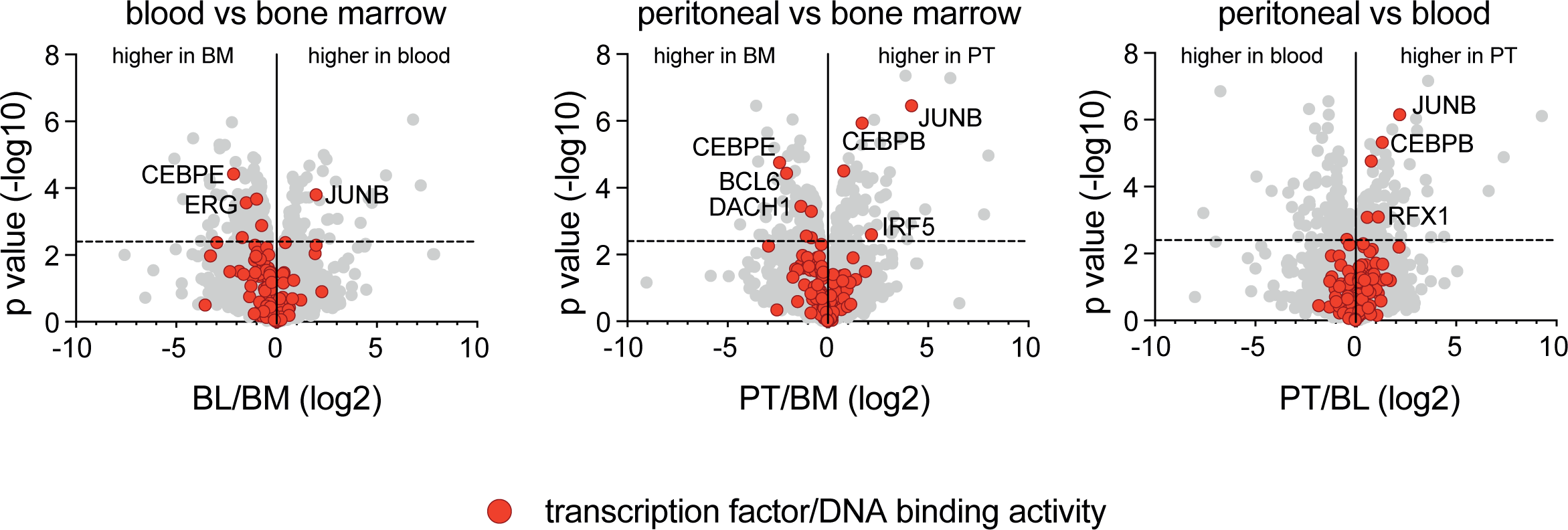
The expression profile of transcription factors in mouse neutrophil populations. Volcano plots show the expression profile of proteins annotated as having DNA binding or transcription factor activity (Gene Ontology term 0003700). For each population 4 biological replicates were generated. BM: bone marrow, BL: blood, PT: peritoneal cavity. The horizontal dashed line on volcano plots indicates q = 0.05.

**Supplementary Figure 4.**
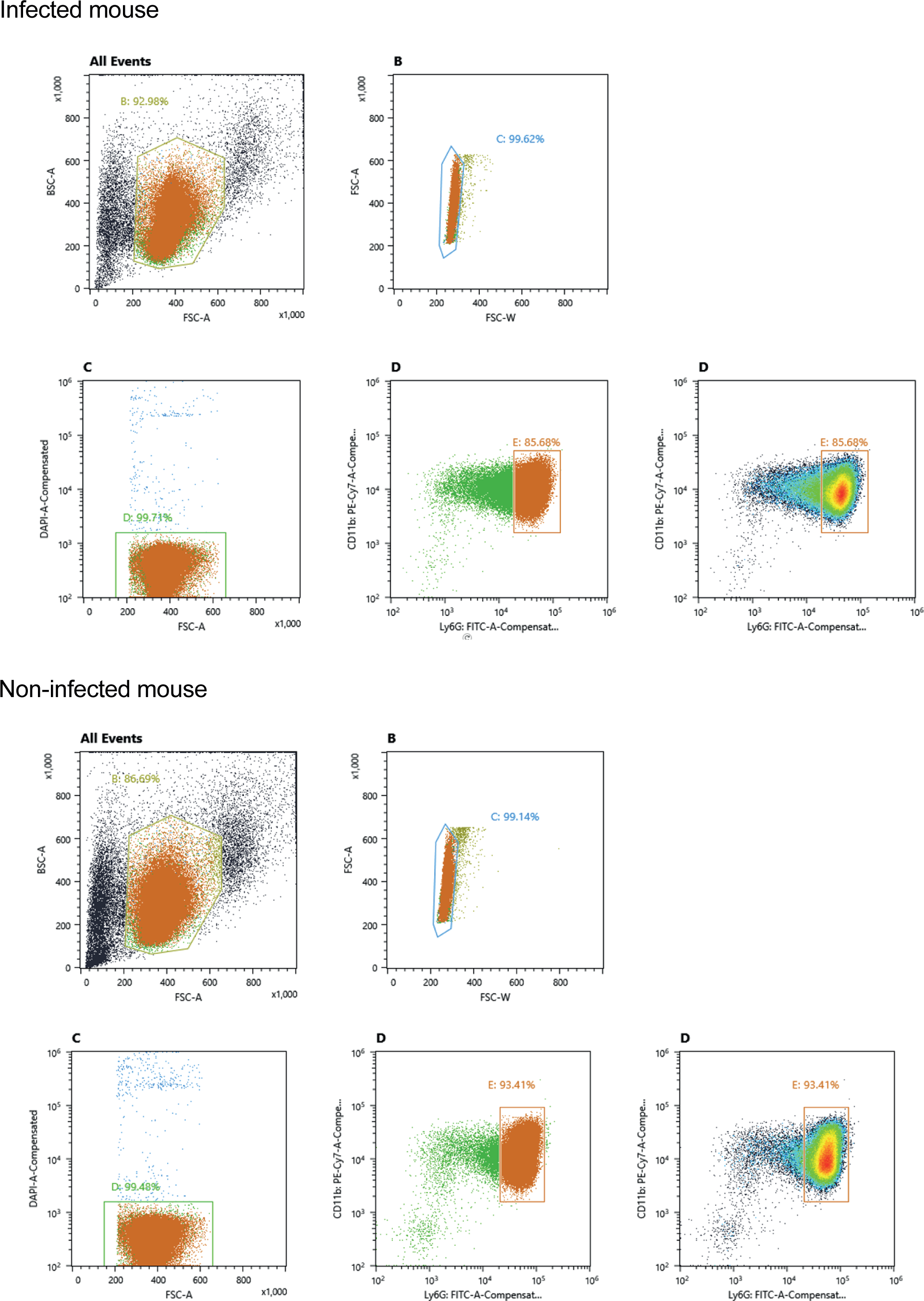
Gating strategy for fluorescent activated cell sorting pure neutrophil populations from *Candida albicans* infection experiment. Neutrophils from infected or control mice (PBS treated) were sorted using Ly-6G FITC, CD11b PE-Cy7 and DAPI.

**Supplementary Figure 5.**
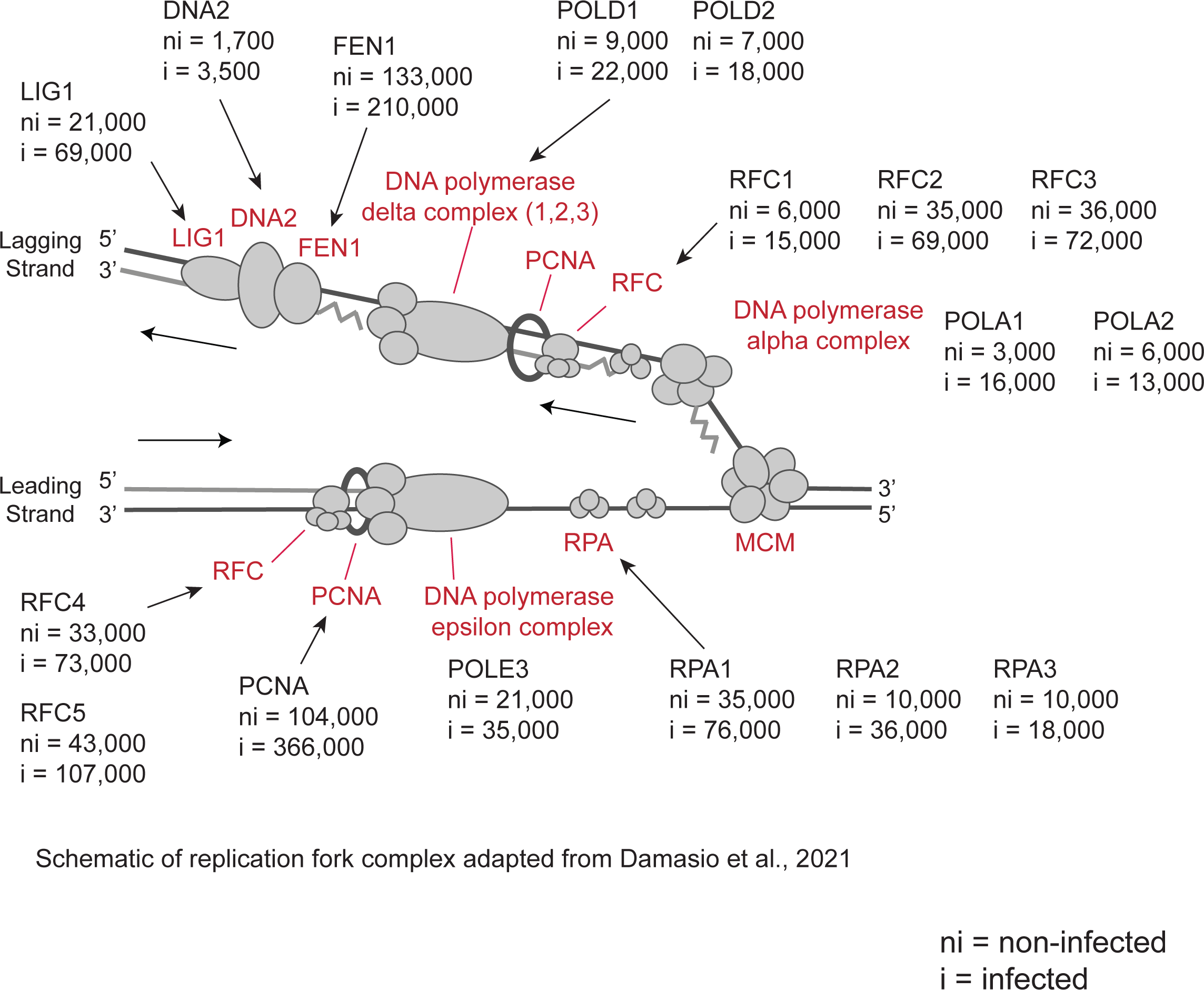
The impact of systemic infection on bone marrow neutrophil DNA replication machinery. Schematic showing the DNA replication fork complex and the average protein copy numbers per cell for infected and non-infected neutrophils. Copy numbers are rounded to the nearest thousand except DNA2 which is rounded to the nearest hundred. For each population 3 biological replicates were generated. Figure adapted from (Damasio et al. 2021)

